# HIV-DRLink: A tool for detecting linked HIV-1 drug resistance mutations in next generation sequencing data

**DOI:** 10.1101/866715

**Authors:** Wei Shao, Valerie F. Boltz, Junko Hattori, Michael J. Bale, Frank Maldarelli, John M. Coffin, Mary F. Kearney

**Affiliations:** Advanced Biomedical Computing Science, Frederick National Laboratory for Cancer Research (FNLCR) sponsored by the National Cancer Institute, Frederick, MD; HIV Dynamics and Replication Program, CCR, National Cancer Institute, Frederick, MD; Department of Molecular Biology and Microbiology, Tufts University, Boston, MA

**Keywords:** HIV, HIV-DRLink, linked drug resistance mutations, Stanford HIVdb, next-generation sequencing, uSGS

## Abstract

The prevalence of HIV-1 drug resistance is increasing worldwide and monitoring its emergence is important for the successful management of populations receiving combination antiretroviral therapy (cART). Using Ultrasensitive Single-Genome Sequencing (uSGS), a next-generation method that avoids PCR bias and PCR recombination, a recent report showed that pre-existing dual-class drug resistance mutations linked on the same viral genomes were predictive of treatment failure while unlinked mutations were not. Because of the large numbers of sequences generated by uSGS and other next-generation sequencing methods, it is difficult to assess each sequence individually for linked resistance mutations. Several software/programs exist to report the frequencies of individual mutations in large datasets but they provide no information on their linkage. Here, we report the HIV-DRLink program, a research tool that provides mutation frequencies in the total dataset as well as their linkage to other mutations conferring resistance to the same or different drug classes. The HIV-DRLink program should only be used on datasets generated by methods that eliminate artifacts due to PCR recombination, for example, standard Single-Genome Sequencing (SGS) or Ultrasensitive Single-Genome Sequencing (uSGS). HIV-DRLink is exclusively a research tool and is not intended to inform clinical decisions.

## 1. Introduction

The advent of combination antiretroviral therapy (cART) changed HIV/AIDS from a deadly disease in most individuals to one that can be managed with lifelong treatment (Jones et al., 2014). However, due to the high genetic diversity of HIV-1, low-frequency drug resistance mutations exist in patients even prior to ART initiation (Coffin, 1995; Gupta et al., 2018; Sapozhnikov et al., 2017) and these low frequency mutations may lead to the emergence of drug resistant viral rebound and treatment failure. Recently, acquired and transmitted drug resistance has become quite prevalent in some parts of the world (Phillips et al., 2014; WHO, 2019) constituting a major barrier to successful treatment of HIV-1.

The Stanford HIV Database (Stanford HIVdb) is a reliable and accurate tool for interpreting HIV drug resistance in population genotypes (Tang, Liu, and Shafer, 2012) (https://hivdb.stanford.edu/). While important, the detection of pre-existing single drug resistance mutations misses the significance of resistance mutations that are linked on the same viral genomes and that were recently shown to be associated with cART failure (Boltz et al., 2019). In fact, detection of pre-existing, linked, dual-class resistance mutations is a better predictor of cART failure than detection of pre-existing, single resistance mutations (Boltz et al., 2019). One difficulty in the interpretation of linked mutations is that bulk PCR may result in artifactual recombination (Shao et al., 2013). Therefore, detection of linked mutations can only be applied to sequences obtained using methods that reduce PCR recombination in conjunction with pipelines that omit sequences resulting from PCR recombination, such as Single-Genome Sequencing (SGS) (Palmer et al., 2005), ultrasensitive Single-Genome Sequencing (uSGS) (Boltz et al., 2016), or other similar next-generation sequencing (NGS) methods (Jabara et al., 2011; Zhou et al., 2015). In brief, uSGS uses primer IDs like many other NGS approaches (Casbon et al., 2011; Fu et al., 2011; Kinde et al., 2011; Liang et al., 2014), however, the Illumina adapters are added by ligation rather than by PCR, significantly reducing the bias and recombination that is inherent to amplification with long PCR primers. Although uSGS can only obtain sequence reads up to about 500 base pairs from amplicons up to 1kb in length, it is possible to link protease inhibitor (PI) resistance mutations to reverse transcriptase (RT) mutations by obtaining 250 base pair reads from one end of the 1kb amplicon and 250 base pair reads from the other end, for example. Significant new changes to the Pacbio platform and chemistry allow for much longer sequence reads (up to the full-length HIV genome) with much higher accuracy than previously possible with this approach (Wenger et al., 2019). Pacbio technology may allow for future investigations of linkage between PI, RT, integrase (IN), and even Env drug resistance mutations (Huang et al., 2016; Van Duyne et al., 2019) While many programs exist for the analysis of HIV sequencing data including their assembly, base-calling, and mutation frequencies (Verbist et al., 2015; Yang et al., 2013; Zagordi et al., 2011) (Brumme and Poon, 2017; Houssaini et al., 2013; Huber et al., 2017; Noguera-Julian et al., 2017; SahBandar et al., 2017), these programs do not detect linkage of drug resistance mutations on the same viral genomes.

Here, we describe a tool called HIV-DRLink that can quickly process thousands of HIV-1 sequences using the Stanford HIVdb server to report, not only the frequencies of single drug resistance mutations in the population, but also the frequency of mutations linked on the same viral genomes, which may be predictive of cART failure. HIV-DRLink is intended as a research tool to efficiently analyze large scale NGS datasets generated specifically using methods that preclude PCR-based recombination. HIV-DRLink is not intended for informing clinical decisions.

## 2. Materials and Methods

### 2.1 HIVDR-Link description

HIV-DRLink is based on the Stanford HIVdb genotypic resistance interpretation program using the Stanford command line program “Sierra Web Service 2.0: 2016 – present” (https://hivdb.stanford.edu/page/webservice/). Therefore, the first step in the HIVDR-Link pipeline is submission of the dataset, in a fasta format, to the Stanford HIVdb. Alignment of sequences and specific information in the sequence headers are not required. For large-scale sequence data, users must download and install a Python client SierraPy from Stanford HIVdb (https://hivdb.stanford.edu/page/webservice/). The output of SierraPy is a JavaScript Object Notation (JSON) file. HIV-DRLink is then used to parse the output file to calculate the frequencies of linked and individual drug resistance mutations in a sample population.

The output of HIVdb Sierra JSON files can be extensive. To simplify the output, a GraphQL protocol is used to select only the gene names (PR, RT, or IN), the mutation types (primary, for example), and the specific mutations. The GraphQL protocol used in the pipeline is a simple text file called “simple_mutations.gql” which is:

~~~
inputSequence {
  header,
},
mutations {
 gene {name}
 primaryType
 text
}
~~~

Two steps are used to run the pipeline:

Step 1: Submit fasta formatted sequences to Stanford HIVdb using the following command line after the Python client SierraPy is installed locally:

~~~
sierrapy fasta input_file.fasta simple_mutations.gql -o output. Json
~~~

Where output.json is an example output file with any name.

Step 2: Run HIV-DRLink.pl on the output file, output.json, from step 1 using the following command line:

~~~
HIV-DRLink.pl output.json
~~~

It should be noted that mutations or polymorphisms in non-drug resistance positions are ignored, and thus the sequences with the same patterns of drug resistance mutations may not be identical at non-resistance sites. The results of HIV-DRLink are reported in a tab delimited text file. HIV-DRLink.pl is available at the GitHub code repository at https://github.com/Wei-Shao/HIV-DRLink.

### 2.2 Meta sequence data

The pipeline was tested for speed and performance using HIV-1 subtype B sequences from Los Alamos HIV Sequence Database (https://www.hiv.lanl.gov/content/sequence/HIV/mainpage.html), including 500 *pol* (RT only) sequences and 200 full-length *pol* sequences (nucleotide positions 2045 to 5200). Because the sequences from the Los Alamos Database typically result from bulk sequencing, not single-genome sequencing, they are used for proof of principle only and, thus, reported mutations may not actually be genetically linked on the same viral genomes. For the pipeline speed and performance test, 23,781 sequences of the RT encoding fragment of *pol* from all subtypes were downloaded from Los Alamos HIV Sequence Database.

### 2.3 Clinical sequence data

While HIV-DRLink can be used to detect linkage of drug resistance mutations in sequences obtained by technologies that omit artifactual recombination, Illumina MiSeq-based uSGS data was used here to validate the pipeline on sequences obtained from a clinical sample (Boltz et al., 2016). The clinically-derived sequences were obtained from genbank (accession #’s KY810858-KY812454). After filtering to remove low quality reads, the paired end fastq files were used for bioinformatics processing to generate HIV-1 sequences of 404 bases in length that covered RT from codons 59 to 131 and 166 to 226 (Boltz et al., 2016).

## 3. Results

### 3.1 Testing the accuracy of HIVDR-Link on sequences obtained from Los Alamos HIV Database

Table 1 shows the results of an HIV-DRLink run on 500 patient-derived HIV sequences obtained from the Los Alamos database. As stated in the methods, although the training dataset contains sequences generated by bulk PCR and sequencing and, therefore, true linkage cannot be determined with such data, it is used here to assess the ability of the pipeline to accurately report drug resistance mutations and to assess its rate of processing large datasets.

**Table 1.**
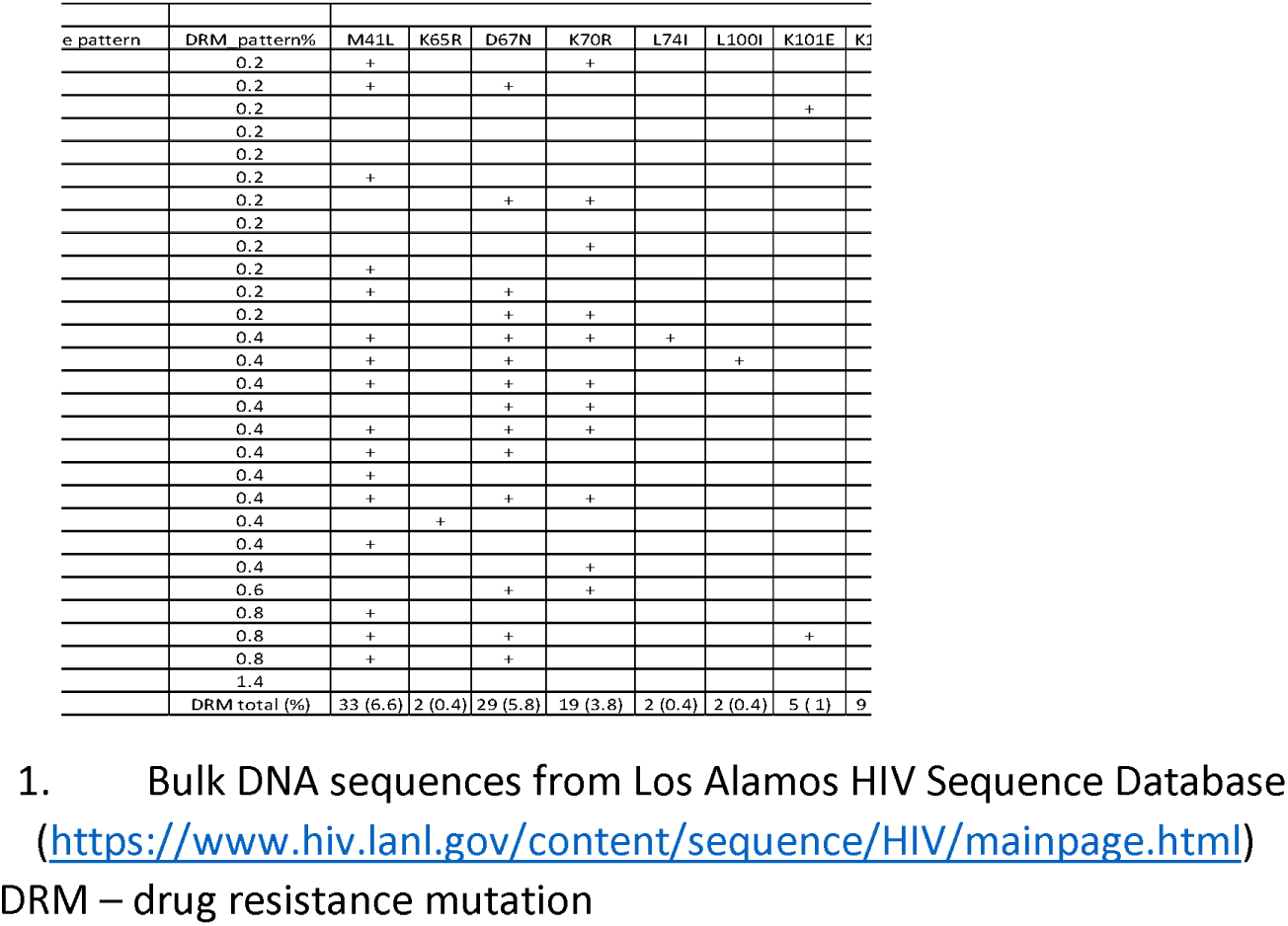
Drug resistance frequencies of HIV-1 subtype B RT sequences^1^.

The first column in Table 1 shows the number assigned to each drug resistance pattern; the second column shows the specific pattern identified, and the third column shows the number of sequence variants that share that particular pattern. Among the 500 sequences retrieved from the Los Alamos HIV Database, 56 had at least one drug resistance mutation (Table 1, bottom row). While some sequences had a single resistance mutation, for example, pattern #3 had only K101E, some others had two or more resistance-conferring mutations, for example, pattern #15 had M41L, D67N, K70R, M184V, L210W, T215F, and K219Q. The percentages of each resistance pattern in the population are shown in the fourth column, ranging from 0.2% to 1.4%. The remainder of the columns show the presence of each individual drug resistance mutation with the last row providing the frequency of each in the total population.

While Table 1 shows the drug resistance patterns in RT only, our program can reveal linkage of mutations in protease (PR) and integrase (IN) and other genes without additional input options or parameters. In addition to the 500 sequences described above, we downloaded an additional 200 bulk *pro-pol* sequences from the Los Alamos Database to test in the pipeline for analysis of linked mutations in PR-RT-IN. Supplemental Table 1 shows an HIV-DRLink output file demonstrating that some sequences, as in the first set of data, had only one resistance mutation, for example, pattern #1 included only S147R in IN while others had “linked” mutations, such as pattern #6 with M46I and N88D in PR, M41L, Y215Y in RT, and G163K in IN and pattern #34 with V32I, L33F, M46I, I47V, I54L, I84V in PR, M41L, D67N, K70R, M184V, T215F, and K219Q in RT, and G140S, Q148H in IN. As stated above, because the training data used was generated by bulk sequencing, the mutational patterns described do not report true linkage of mutations on single genomes but demonstrate the accuracy of the program to report patterns of drug resistance mutations in a hypothetical large dataset.

### 3.2 Drug resistance detection in clinical samples using HIV-DRLink

To evaluate true linkage of drug resistance mutations on single HIV-1 genomes, we tested the pipeline on plasma HIV-1 RNA sequences obtained by uSGS (Boltz et al., 2016) (sequences available at KY810858-KY812454). The plasma sample was obtained from an HIV-infected donor with viremic failure on ART. uSGS yielded 1597 high quality single genome *pol* sequences covering RT codons 59 to 131 and 166 to 226 (Boltz et al., 2016). Table 2 shows 12 different resistance patterns that were detected using the HIV-DRLink program, all of which had linked mutations. While some of the patterns were rare, for example, #1 to 3 comprising 0.06% of the population, pattern #12 with 4 linked mutations comprised 73% of the population. HIV-DRLink also calculated the levels of individual drug resistance mutations in the sample. For example, 21.54% of the sequences had the D67N mutation and 99% had T215Y.

**Table 2.** Frequencies of linked drug resistance in HIV1 subtype B RT sequences^1^.

| DRM pattern # | DRM pattern | # sequence with same pattern | RT DRM |  |  |  |  |  |  |  |
| --- | --- | --- | --- | --- | --- | --- | --- | --- | --- | --- |
|  |  |  | DRM pattern% | D67N | K101E | Y188C | G190A | L210W | T215F |  |
| 1 | L210W,T215Y | 1 | 0.06 |  |  |  |  | + |  |  |
| 2 | K101E,G190A,L210W,T215F | 1 | 0.06 |  | + |  | + | + | + |  |
| 3 | K101E,Y188C,G190A,L210W,T215Y | 1 | 0.06 |  | + | + | + | + |  |  |
| 4 | D67N,K101E,G190A,L210W | 2 | 0.13 | + | + |  | + | + |  |  |
| 5 | K101E,L210W,T215Y | 2 | 0.13 |  | + |  |  | + |  |  |
| 6 | G190A,L210W | 2 | 0.13 |  |  |  | + | + |  |  |
| 7 | K101E,G190A,T215Y | 2 | 0.13 |  | + |  | + |  |  |  |
| 8 | K101E,G190A,L210W | 11 | 0.69 |  | + |  | + | + |  |  |
| 9 | D67N,G190A,L210W,T215Y | 16 | 1.00 | + |  |  | + | + |  |  |
| 10 | G190A,L210W,T215Y | 60 | 3.76 |  |  |  | + | + |  |  |
| 11 | D67N,K101E,G190A,L210W,T215Y | 326 | 20.41 | + | + |  | + | + |  |  |
| 12 | K101E,G190A,L210W,T215Y | 1173 | 73.45 |  | + |  | + | + |  |  |
| Summary | Total (% based on 1597 total input seq) | 1597 (100) | DRM total (%) | 344 (21.5) | 1518 (95.1) | 1 (0.06) | 1594 (99.8) | 1595 (99.9) | 1 (0.06) | 15 |

1. Clinical sequences analyzed by uSGS (Boltz et al., 2016)
DRM – drug resistance mutations

### 3.3 Speed of HIV-DRLink

To evaluate HIV-DRLink speed and performance, 23,781 *pol* sequences from the Los Alamos HIV Sequence Database were submitted to the Stanford HIVdb via Python client Sierrapy (https://hivdb.stanford.edu/page/webservice/). While it took approximately 40 minutes to obtain a list of mutations for each sequence from the Stanford Database using Python client SierraPy, it took approximately 1 minute to extract the data and produce the final result using HIV-DRLink.

## 4. Discussion

We developed and applied a program called HIV-DRLink that is capable of reporting linked and unlinked HIV-1 drug resistance mutations in large-scale NGS datasets generated by uSGS (Boltz et al., 2016) or other similar methods (Jabara et al., 2011), making use of the very well annotated and maintained Stanford HIV Drug Resistance Database (https://hivdb.stanford.edu/). Although other programs that calculate mutation frequencies from such data are available (Houssaini et al., 2013; SahBandar et al., 2017), they only calculate the frequencies of individual resistance mutations in the HIV-1 population and do not report their linkage. Mutations linked on the same HIV-1 genomes have been shown to be associated with virologic failure even when present at the same or lower levels as single mutations (Boltz et al., 2019) and, therefore, are important to detect and report for investigational use. Of importance, HIV-DRLink was developed for the detection of linked mutations from NGS data generated by methods such as uSGS (Boltz et al., 2016) and other similar methods (Jabara et al., 2011; Zhou et al., 2015) where PCR recombination has been either eliminated, as is the case with uSGS, or shown to be minimal. HIV-DRLink output from data obtained using other NGS methods must be interpreted with care since linkage of mutations may be lost or generated by PCR-based recombination.

To test the accuracy and speed of HIV-DRLink, HIV-1 *pol* sequences from the Los Alamos HIV Sequence Database (http://www.hiv.lanl.gov/) were queried. The results show that HIV-DRLink accurately and rapidly reported the frequency and linkage of individual drug resistance mutations. It should be noted that the sequences from the Los Alamos HIV Database are mostly from population sequencing and each represents a mixture of genomes so that putatively linked mutations may, in fact, be on separate molecules. HIV-DRLink was further tested on uSGS data obtained from a clinical specimen where each sequence was known to have originated from a single viral template (Boltz et al., 2016) and the pipeline reported accurate results in less than one minute. In conclusion, we developed a tool, HIV-DRLink, to rapidly detect linked and unlinked HIV-1 DRMs in large scale NGS datasets.

## Acknowledgements

We acknowledge that a very important part of the pipeline is to obtain information from Stanford HIVdb (https://hivdb.stanford.edu/). We thank Mr. Philip Tzou and Dr. Robert Shafer of Stanford HIV Drug Resistance Database for very valuable discussions. We thank Connie Kinna, Anne Arthur, Valerie Turnquist, and Sue Toms for administrative support. We acknowledge the funding sources for this study from NCI CCR, the Office of AIDS Research, NIH, NCI intramural funding to MFK, and NCI Contract No. HHSN261200800001E. JMC was a Research Professor of the American Cancer Society and was supported by Leidos Contract 13XS110.

